# Bacterial communication and relevance of quantum theory

**DOI:** 10.1101/145458

**Authors:** Sarangam Majumdar, Sisir Roy

## Abstract

The recent findings confirm that bacteria communicate each other through chemical and electrical signals. Bacteria use chemical signaling molecules which are called as quorum sensing molecules(QSMs) or autoinducers. Moreover, the ion channels in bacteria conduct a long-range electrical signaling within biofilm communities through propagated waves of potassium ions and biofilms attracts other bacterial species too. Both communication process are used by bacteria to make their own survival strategies. In this article, we model this bacterial communication mechanism by complex Ginzburg- Landau equation and discuss the formation of patterns depending on kinematic viscosity associated with internal noise. Again, the potassium wave propagation is described by the non-linear Schr*ö*dinger equation in a dissipative environment. By adding perturbation to non-linear Schr*ö*dinger equation one arrives at Complex Ginzburg-Landau equation. In this paper we emphasize that at the cellular level(bacteria) we use Complex Ginzburg - Landau equation as a perturbed Nonlinear Schr*ö*dinger equation to understand the bacterial communication as well as pattern formation in Biofilms for certain range of kinematic viscosity which can be tested in laboratory experiment. Here, the perturbation is due to the existence of non thermal fluctuations associated to the finite size of the bacteria. It sheds new light on the relevance of quantum formalism in understanding the cell to cell communication.

## I. INTRODUCTION

Bacteria are unicellular microorganism which is found in everyday life. They are attached with human being too. It has been observed that some Gram positive and Gram negative bacteria live in organized communities [1]. This bacterial communities are glued together by extracellular polymeric substance (EPS), which is called a biofilms[2]. Bacteria in biofilm communities can communicate to each other in two different ways. First, bacteria cooperate with each other by releasing different chemical signaling molecules or autoinducers (AI-1 and AI-2). Autoinducers are wide variety of molecules which are used by bacteria for inter- species and intra-species communication process. As the bacterial colonies are dense, then the signaling molecules are secreted into extracellular locations and accumulated. When a threshold concentration of the signaling molecules is achieved, a coordinated change in bacterial collective behavior is initiated. This bacterial collective behavior is known as quorum sensing by which bacterial cell integrate in order to determine their optimal survival stagey[1, 2]. Quorum sensing coordinate the gene expression, when the bacterial cell population has reached a high cell density. In a Gram negative bacteria acyl homoserine lactone (AHL) and quinolone mediated quorum sensing is already observed[1]. Gram positive bacteria uses autoinducer peptide (AIP), which ranges from 5 to 34 amino acids in length and typically contain unusual chemical architectures. [1]. Quorum sensing is responsible for mediating a variety of social activities in biofilms, which include the swarming motility, biofilm dispersion, biofilm growth and antimicrobial resistance etc. It is found that quorum sensing regulating EPS production during biofilm formation[2].

Bacterial biofilms are very organized communities which contain billions of densely packed bacterial cell. Recent studies show that apart from using chemical molecules, bacteria can communicate over a long distances which is enabled by bacterial ion channels[2].This electrical communication within bacterial communities occur through spatially propagating waves of potassium[2]. Cells insides the biofilm communities can cooperate and compete with each other for resources. When these communities grow larger, the supply of nutrient to the interior cells becomes limited because the nutrient consumption is increasing. At the same time, this nutrient consumption is associated with the growth of multiple layers of cells in the biofilm periphery. This periphery not only protect interior cells from the external attack but also starve them through nutrient consumption. The conflict between starvation and protection is resolved through emergence of long-range metabolic co-dependence between interior and peripheral cells and gives rise to collective oscillations within biofilms [2].This electrically oscillating biofilms attract distant motile cells toward the biofilms[2]. In 2017, it has been discovered that two *B*. *subtilis* biofilm communities undergoing metabolic oscillations become coupled through electrical signaling and synchronize their growth dynamics. This Coupling increases competition by also synchronizing demand for limited nutrients[2]. They confirm that biofilms resolve this conflict by switching from in phase to anti- phase. Different biofilm communities take turns consuming nutrients. Thus distant biofilms can coordinate their behavior to resolve nutrient competition through time-sharing. This is a very intelligent and efficient strategy to share the limited resources [2]. In this presentation, we are briefly discuss the theory.

## II. THEORY

The densely packed bacterial populations develop a coordinated motion on the scales length 10*µm* to 100*µm* in comparison to the size of a each single bacterium of order 3*µm* when the bacterial cell density reaches a sufficiently high value. We assume that the collective behavior of the densely packed bacteria inside the biofilm is similar to the behavior of the of the dense granular systems[3].The dense granular system usually behave like a fluid which is quite different from the ordinary fluids. The finite size of the bacteria indicates the existence of an intermediate length scale, which leads us to introduce a source of fluctuation which is quite different than thermodynamic fluctuation. This new type of fluctuation can be considered as a non-local noise. The swimming induced stress on the bacteria that can change the local arrangement of bacteria induce stress fluctuations. This stress fluctuation can lead to shear motion and hence is called non-local. Thus, two different type of noise are present in the bacterial communication system and dominance of one over the other depends on the force 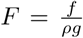 which is applied to the complex biological system where *f* be the volume density of the forcing and *g* is the acceleration due to gravity [3].

### A. Viscosity and non-local theory

Let us consider the state space(*ρ*, **v**) of one component fluid, where *ρ* be the density and **v** be the velocity of the fluid. The stress tensor and/or the pressure term is the only constitutive quantity in this framework. We consider the higher order derivatives of the basic variable (density and velocity) to extend theory of weakly non-local hydrodynamics. Without loss of generality, the balance of mass and momentum can be expressed as

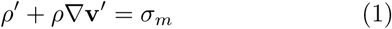

and

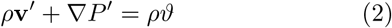

Here *P* is the pressure and *ϑ* be the force density. This is formally known as Cauchy momentum equation. Now, we can extend this framework by considering the state space spanned by (*ρ*, ∇*ρ*, **v**, ∇**v**, ∇^2^*ρ*).

Van and Fulop [4] show that there exists a scalar valued function *ϕ*_*v*_ or non-local potential such that

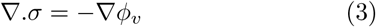

where *ϕ*_*v*_ is the course- grained potential or kinematic viscosity potential and *σ*_*ij*_ be shear tensor. One can calculate the viscosity potential from the entropy density function

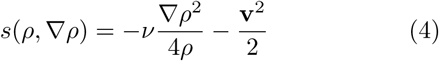

The non-local potential can be written as

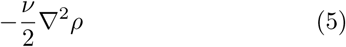

where *ν* is kinematic viscosity and 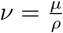 (*μ* is dynamical viscosity of the fluid). We define a kinematic velocity as 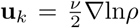 [5], which is depend upon the cell density. Here we introduce the kinematic velocity in order to relate to a kind of fluctuations due to the existence of finite length scale associated to granular nature of the fluid. Finally (after some algebraic calculation), we get a general expression as

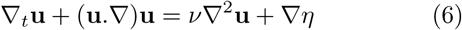

where ∇*η* = −*ν*∇^2^(∆**u**_*k*_) and ∆**u**_*k*_ = **u** – **u**_*k*_. The above Eq.(6) is known as noisy Burgers equation. We emphasize that the non-local hydrodynamical model (based on Ginzburg-Landau framework) can explain the quorum sensing phenomena in a consistent way. The rearrangement of the configuration of the coarse grained systems (bacterial rearrangement within biofilms communities) produce a noise which give rise to kinematic viscosity. This noise induces kinematic viscosity which plays a crucial role to understand the quorum sensing phenomena. This mathematical framework gives the view of an internal structure of the complex biological communication system and viscosity is the property which makes the bacterial cells stick together into clusters predicted by Zel-dovich approximation, just mimicking gravitational effect on the smaller scales [3].This approximation can describe the general structure of this nonlinear biological phenomena. It is to be mentioned that the origin of viscosity is associated with the weakly non-local effects in the internal structure of the system. One of the present authors (SR) along with Llinas [3] showed that kinematic viscosity plays a vital role in forming the metastable states of the bacteria responsible for quorum sensing. It is interesting to note that various type of patterns are formed by bacteria in Biofilm. Now we study the formation of patterns in Biofilms and the role of kinematic viscosity.

### B. Kwak Transformation and Reaction- diffusion systems

The quorum sensing system is modeled by noisy Burger equation (Eq.6). We can rewrite the Eq.(6) as

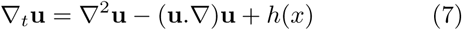

with 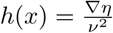.

By using Kwak transformation 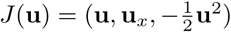 we can obtain a new system as [6] (see Appendix for detail calculation and properties)

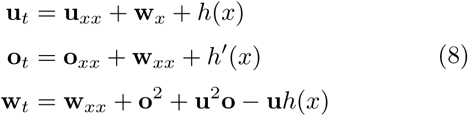

The above Eq.(8) is a reaction- diffusion system which gives the mathematical framework for the pattern formation.

### C. Pattern Formation and Viscosity

In this multicellular system bacterial cells form a different patterns based on chemical gradients of QSM signal that is synthesized by quorum sensing bacterial cells. The above theoretical analysis reveal that parameters (kinematic viscosity and noise) most significantly affect to form patterns over space and time. Furthermore, the mathematical approach is able to predict how the system behaves if we change the initial value. We can say that these are crucial parameters (kinematic viscosity and noise) of the system. It should be noted that the regulatory behaviors mentioned above are nontrivial consequence of the model. In our system, we observed that the quorum takes place in a certain range of kinematic viscosity [0.01, 0.32]*m*^2^/*s* which is considered as very small viscosity of the fluid (see detail in [6]). We also used different numer- ical scheme and initial data to show the the quorum sensing system behaviour. The behaviour changes with the initial data and system forms different wave patterns.

### D. Electrical Communication and Non-linear Schr*ö*dinger equation

The recent findings suggest that bacteria communicate through electrical signalling using waves associated to Potassium ions. One of the present authors (SR) along with Rodolfo Llinas showed that Potassium ions follow non-linear Schrodinger equation [7]. This equation can be written in the following form:

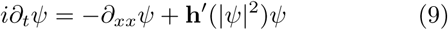

where *ψ* is the wave function of Potassium ion. Now one arrives Complex Ginzburg-Landau equation by adding perturbation to the above non-linear Schr*ö*dinger equation [8]. A Melnikov approach

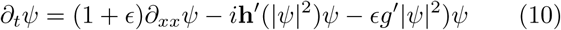

Here *ψ(x,t)* is a complex field and *ϵ* > 0 while **h** = **h**(Ξ) and *g* = *g*(Ξ) are real analytic functions over [0, inf). This non-linear Schr*ö*dinger equation is valid at the level of ion channel where as the perturbation becomes predominant at the cellular level. At the cellular level, the non-thermal fluctuation arises due to the presence of finite size of the cell or grain the granular medium. This fluctuation gives rise to the perturbation on nonlinear Schr*ö*dinger equation and we get generalized Complex Ginzburg-Landau(GL) equation. This Complex GL equation is used for the description of cellular communication through the chemical molecules and also needed to understand the generation of various patterns in Biofilms.

## III. RESULTS AND DISCUSSION

It is evident from the above analysis that we are dealing with two level system in understanding the cellular communication: at the level of ion channel one needs non-linear Schr*ö*dinger equation to understand the communication through electrical signaling through Potassium waves and at the level of chemical signaling one needs Complex Ginzburg-Landau equation. This Complex Ginzburg-Landau equation arises due to the perturbation of non-linear Schr*ö*dinger equation arises due to the existence of non thermal fluctuation. This perturbation in non-linear Schr*ö*dinger equation due to nonthermal fluctuation will be studied in details by considering the effect of noise in such non-linear Schr*ö*dinger equation in subsequent publication. The pattern formation in Biofilms can be tested in the laboratory for the specified range of viscosity.

## IV. APPENDIX-I

Presently, we address the steady state solution of Eq.(7) and Eq.(8). Consider solution of Eq.(7) finite and with a unique steady state solution for small force. Our proposed model for the quorum sensing mechanism 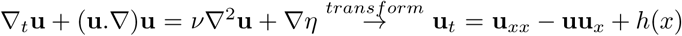 by letting 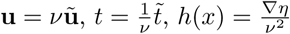. so that the viscosity appears in the noise term.

Mean value of **u** is

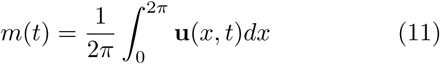

Rate of change of *m* w.r.t time

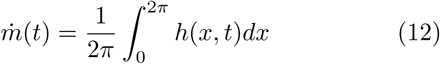

The term *h* will be assumed to have zero mean so that by Eq.(8) the mean of concentration of QSM **u**(*x*,*t*) is conserved.

The solution of the Eq.(7) is treated as a solution of the reaction diffusion system by introducing a nonlinear change of variables. Let **u** be the solution of Eq.(7) and 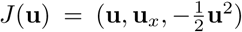. Then (**u**, **o**, **w**) = *J*(**u**) satisfy Eq.(7). The mean of **u** in Eq.(8) is conserved since **u** has zero mean and the mean of **o** is also conserved if **u** satisfy the periodic boundary condition. The mean of **w** is not conserved [6].

To conserve the mean of **w**, we modify Eq.(8) with

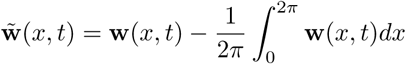

such that the drift free reaction-diffusion system becomes

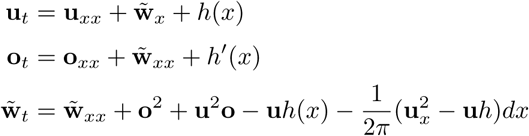

### REMARK

1. If **o**(*x*, 0) = **u**_*x*_(*x*, 0) then **o**(*x*, *t*) = **u**_*x*_(*x*, *t*) ∀ *t* ≥ 0
2. For any steady state solution of Eq.(8) **u***_x_ =* **o**.
3. For any steady state solution of Eq.(8) 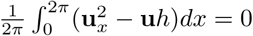
4. Let 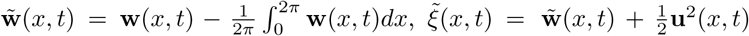 and 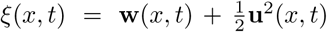, Then 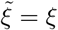 as steady state.

The steady state solution of the proposed model for QS system

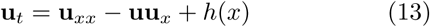

is also the steady state solution to the transformed reaction-diffusion system of the Burger’s equation.

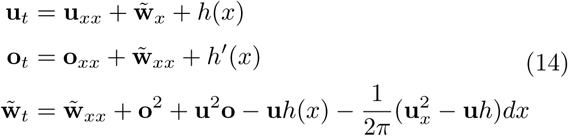

Conversely, any steady state solution 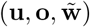 of Eq.(12) is necessarily of the form 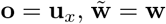 with 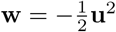 and **u** is the steady state solution of Eq.(11).

### REMARK

1. Every solution to the noisy Burger’s equation (**u**_*t*_ = **u**_*xx*_ – **uu**_*x*_ + *h*(*x*)) satisfies the inequality 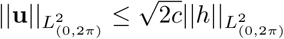 for *t* ≥ *t*_0_ with

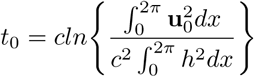

and *c* is the Poincare constant.
2. The steady state solution **u** of forced Burger’s equation satisfies the following inequalities

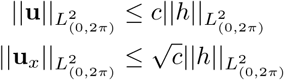
3. There is a unique steady state solution to the noisy Burger’s equation (**u**_*t*_ = **u**_*xx*_ – **uu**_*x*_ + *h*(*x*)). When *h* satisfies

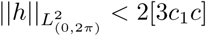

where *c* is the Poincare constant and *c*_1_ is the Sobolev constant.

## V. APPENDIX-I

The reaction -diffusion system can be expressed as

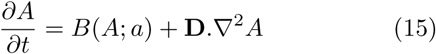

where *A*(*x, t*) is the concentration of quorum sensing molecules, which is depend on space and time and **D** diffusion matrix. We can write the solution in the form as

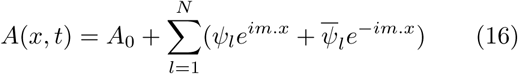

This reaction -diffusion can be described by the complex Ginzburg-Landau equation.

From complex Ginzburg-Landau equation we get,

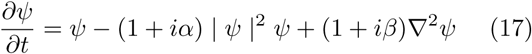

It can also be reduced to a dissipative extension of the nonlinear Schr*ö*inger equation in the form [5]

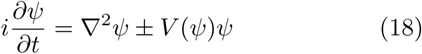

and it admits the solution which can be representation as

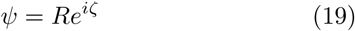

## References

[1] Majumdar S, Mondal S (2016) Conversation game: talking bacteria. J Cell Commun Signal 10(4):331–335

[2] Majumdar S, Pal S (2017) Bacterial intelligence: imitation games, time-sharing, and long-range quantum coherence. J. Cell Commun Signal. Doi: 10.1007/s12079-017-0394-6

[3] Roy S, Llinas R (2016) Non-local hydrodynamics of swimming bacteria and self-activated process. BIOMAT 2015 Proceedings of the International Symposium on Mathematical and Computational Biology. World Scientific. 153–165.

[4] Van P, Fulop T, (2004) Weakly Nonlocal Fluid Mechanics The Shro¨dinger Equation, arXiv:quant-ph/0304062v2

[5] A. L. B. Ribeiro and J. G. Peixotode Faria, (2005) Weakly Non local Hydrodynamics and The origin of Viscosity in the Adhesion Model. Physical Review D 71, 067302

[6] Majumdar S, Roy S, Llinas R (2017) Bacterial Conversations and Pattern Formation. bioRxiv. doi: 10.1101/098053

[7] Roy S, Llins R (2009) Relevance of quantum mechanics on some aspects of ion channel function. Comptes rendus biologies 332(6): 517–522

[8] Gustavo Cruz-Pachecoa, C.David Levermoreb, Benjamin P. Lucec(2004) Complex GinzburgLandau equations as perturbations of nonlinear Schr*ö*dinger equations; Physica D 197, 269–285.

